# The reovirus σ3 protein inhibits NF-κB-dependent antiviral signaling

**DOI:** 10.1101/2021.10.05.463132

**Authors:** Andrew J. McNamara, Austin D. Brooks, Pranav Danthi

**Author notes:** To whom correspondence should be addressed: Department of Biology, Indiana University, Bloomington, IN 47405.

## Abstract

Viral antagonism of innate immune pathways is a common mechanism by which viruses evade immune surveillance. Infection of host cells with reovirus leads to the blockade of NF-κB, a key transcriptional regulator of the hosts’ innate immune response. One mechanism by which reovirus infection results in inhibition of NF-κB is through a diminishment in levels of upstream activators, IKKβ and NEMO. Here, we demonstrate a second, distinct mechanism by which reovirus blocks NF-κB. We report that expression of a single viral protein, σ3, is sufficient to inhibit expression of NF-κB target genes. Further, σ3-mediated blockade of NF-κB occurs without changes to IKK levels or activity. Expression of only a subset of NF-κB target genes is reduced. Among NF-κB targets, the expression of type I interferon is significantly diminished by σ3 expression. Correspondingly, ectopic expression of σ3 enhances viral replication. Expression of NF-κB target genes varies following infection with closely related reovirus strains. Our genetic analysis identifies that these differences are controlled by polymorphisms in the amino acid sequence of σ3. This work identifies a new role for reovirus σ3 as a viral antagonist of the NF-κB-dependent antiviral pathways.

**IMPORTANCE:** Host cells mount a response to curb virus replication in infected cells and prevent spread of virus to neighboring, as yet uninfected cells. The NF-κB family of proteins is important for the cell to mediate this response. In this study, we show that a single protein, σ3, produced by mammalian reovirus, impairs the function of NF-κB. We demonstrate that by blocking NF-κB, σ3 diminishes the hosts’ response to infection and promotes viral replication. This work identifies a second, previously unknown mechanism by which reovirus blocks this aspect of the host cell response.

## INTRODUCTION

In order to successfully infect a mammalian host, viruses need to antagonize the innate immune response (1, 2). The innate immune response in mammals is initiated by a sensor, which leads to the activation of multiple antiviral transcription factors. RNA viruses can be sensed extracellularly or within endosomes via Toll-like Receptors (TLRs). Alternatively, RNA viruses can be sensed in the cytoplasm by RIG-I-like Receptors (RLRs) (3). Engagement of TLRs or RLRs lead to the activation of transcription factors, which induce expression of antiviral effector proteins (3). These proteins can either antagonize the viral replication cycle in the initially infected cell, or can signal to neighboring cells via production of cytokines including interferons (IFNs), to prevent the establishment of infection (1). The two primary transcription factors which drive the innate immune response are Nuclear Factor-κB (NF-κB) and Interferon Regulatory Factor 3 (IRF3) (3). These two transcription factors, acting either alone or in combination, control the expression of hundreds of target genes. Because these two transcription factors regulate such a wide variety of antiviral factors, they are frequent targets of viral antagonism.

The NF-κB transcription factor family is composed of five different subunits that function as homo or heterodimers. The classical NF-κB complex (henceforth referred to as NF-κB), composed of p65 and p50 subunits, is a critical regulator of antiviral gene expression (4). In an inactive state, it is sequestered in the cytoplasm by the Inhibitor of κB (IκB) inhibitor proteins (4). NF-κB activity is regulated by the IκB Kinase (IKK) complex. The IKK complex, which is composed of IKKα and IKKβ kinases, and the NEMO regulatory protein, phosphorylates IκB (4). Phosphorylation of IκB leads to its proteasomal degradation. This exposes the nuclear localization signal on NF-κB, causing nuclear translocation and accumulation of the transcription factor. In addition to the phosphorylation of IκB, the IKK complex also phosphorylates p65, which is required for the transactivation activity of NF-κB (5). Transcription of NF-κB target genes is also affected by the interaction of NF-κB with transcriptional coregulators, the interaction of other transcription factors with regions adjacent to the NF-κB binding sites, and the chromatin status of genomic DNA in that region.

Mammalian orthoreovirus (reovirus) is a dsRNA virus which replicates in the cytoplasm of host cells (6). The reovirus genomic RNA is a potent activator of the innate immune system (7, 8). Sensing of reovirus genomic RNA by RLRs leads to the activation of IRF3 and NF-κB, which lead to the production of interferon (IFN) and other inflammatory cytokines (9-12). (13). Consistent with this, the genetic absence of either IRF3 or NF-κB exacerbates reovirus disease in a newborn mouse model (14, 15). The regulation of NF-κB in reovirus infected cells is complex. NF-κB activity is observed in reovirus infected cells at early stages of infection (16-18). Later following infection, NF-κB-dependent gene expression is no longer active despite the continued presence of viral RNA within the cytoplasm (13, 17). At this stage, addition of exogenous NF-κB agonists such as Tumor Necrosis Factor alpha (TNFα) also fails to stimulate NF-κB, indicating that the function of a signaling component common to both RLR and TNFα signaling is compromised (13, 17). Inhibition of NF-κB occurs through the loss of two components of the IKK complex, IKKβ and NEMO, The loss of IKKβ and NEMO is dependent on viral gene expression (13). We hypothesize that adequate accumulation of one or more viral proteins late in infection contributes to the loss of IKKβ and NEMO, consequently blocking NF-κB.

In this study, we sought to identify the viral factor responsible for inhibition of NF-κB. We discovered that expression of a single reovirus gene product, σ3, is sufficient to inhibit NF-κB-dependent gene expression following treatment with TNFα or viral genomic dsRNA. Remarkably, this inhibition is neither due to the loss of the IKK complex nor due to impairment of IKK activity. σ3 expression only inhibits the transcription of a subset of NF-κB target genes, suggesting that σ3 inhibits a nuclear function of NF-κB. We demonstrate that σ3 expression augments reovirus replication likely via inhibition of antiviral NF-κB targets such as type I IFN. Finally, using σ3 mutant viruses, we show that properties of σ3 display differences in NF-κB signaling and IFN expression in the context of infection. Thus, our work uncovers a new role for σ3 in inhibiting the NF-κB-driven innate immune response and promoting viral infection.

## RESULTS

### σ3 inhibits NF-κB-dependent gene expression

We recently showed that de novo reovirus gene expression is required for blocking the activity of transcription factor NF-κB (13). Based on these results, we surmised that one or more newly synthesized viral gene products inhibit the activity of NF-κB. To determine if a single viral gene product is responsible for NF-κB inhibition, we co-transfected HEK293 cells with vectors expressing each protein from prototype reovirus strain T3D^F^, with a NF-κB-dependent firefly luciferase reporter (Fig. 1A). After treatment with TNFα, a known NF-κB agonist, we measured NF-κB-dependent luciferase production. While most reovirus genes did not significantly affect TNFα-induced luciferase activity, expression of the S4 gene-encoded σ3 protein inhibited NF-κB-driven luciferase activity by more than 90%. These protein expression constructs were left untagged because we could not determine if tagged proteins would remain functional. Since available antisera to detect each of these proteins vary in their detection sensitivity, we cannot rule out the possibility that the absence of effect of some of these proteins on NF-κB activity is related to lower level expression of these proteins. Nonetheless, our data very clearly demonstrate that σ3 expression inhibits NF-κB activity. The studies presented here are only focused on defining how σ3 blocks NF-κB signaling. The steady state levels of σ3 from our expression vector is lower than those observed for σ3 in cells infected with reovirus, suggesting that the observed phenotype is not related to an artifact of unusually high σ3 expression (Fig. 1B). These data suggest that σ3 expression is sufficient to block NF-κB function in reovirus infected cells.

**Fig 1.**
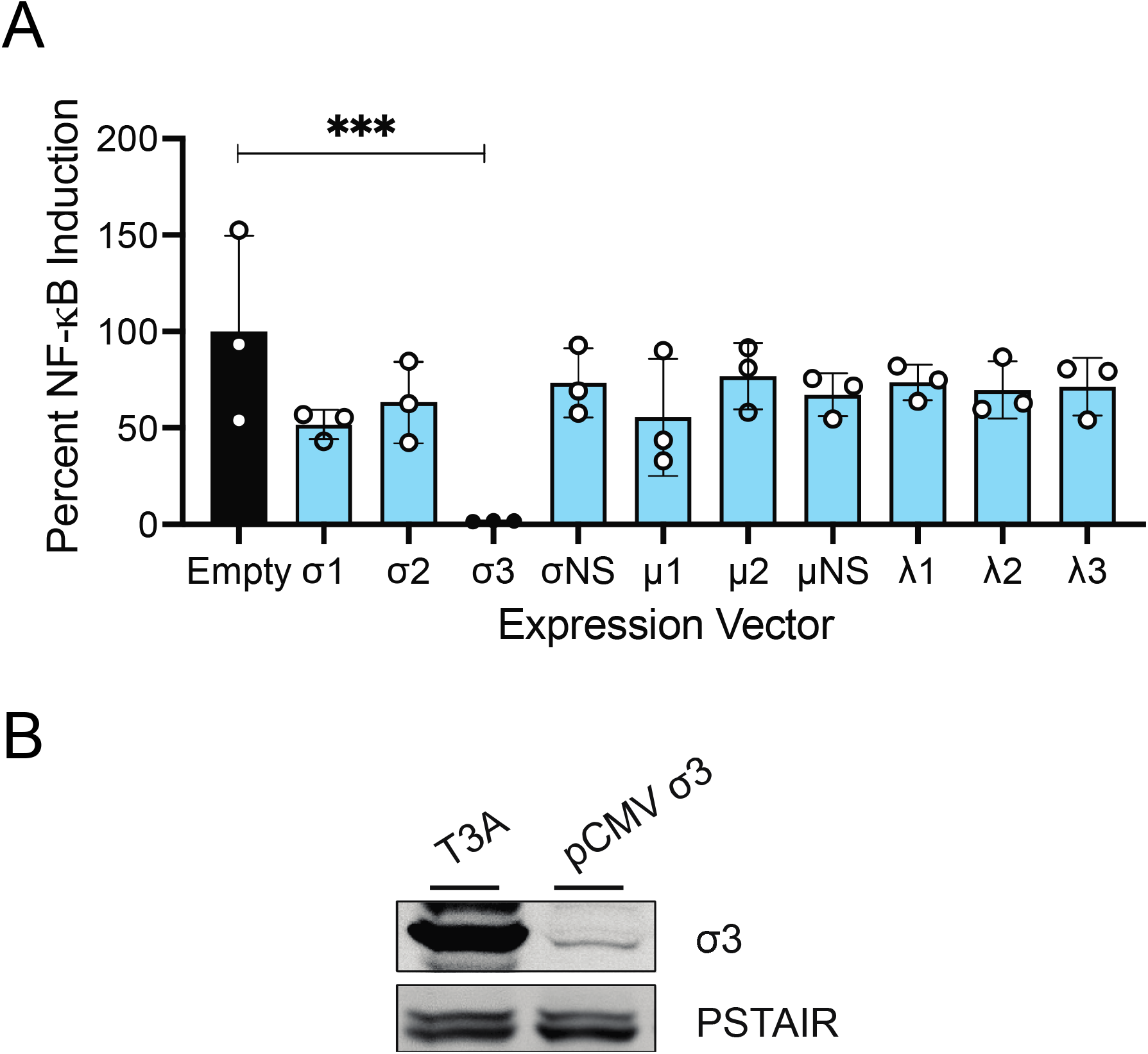
Reovirus protein σ3 inhibits NF-κB-dependent gene expression. (A) HEK293 cells were transfected an empty vector or with vectors corresponding to each reovirus genome segment, an NF-κB-driven Firefly luciferase reporter, and a control Renilla luciferase reporter. Following incubation at 37°C for 24 h, HEK293 cells were treated with 10 ng/ml TNFα. Following additional incubation at 37°C for 7 h, the ratio of Firefly to Renilla luciferase activity (Relative NFκB activity) was quantified. Relative NFκB activity in empty vector transfected cells was set to 100%. Activity for each independent transfection and treatment, the mean value and SD are shown. ***, P < 0.001 by one-way ANOVA with multiple comparisons. (B) HEK293 cells were transfected with an empty vector or a vector expressing σ3. Alternatively, the cells were absorbed with 10 PFU/cell of reovirus strain T3A. Following an additional incubation at 37°C for 24 h, whole cell extracts were immunoblotted using antiserum specific for reovirus and PSTAIR.

### σ3 expression does not affect IKK activity

Our previous work indicated that inhibition of NF-κB-dependent gene expression by reovirus is due to the loss of the IKKβ and NEMO proteins, which comprise the IKK complex that functions upstream of NF-κB (13). Thus, in reovirus-infected cells, even upon addition of NF-κB agonists such as TNFα, NF-κB fails to translocate to the nucleus and its p65 subunit remains unphosphorylated. To determine if σ3 inhibits NF-κB-dependent gene expression due to loss of the IKK complex, we transfected HEK293 cells with a σ3 expression vector and monitored IKK activity by determining the capacity of TNFα to promote NF-κB nuclear translocation. We found that TNFα-driven nuclear accumulation of p65 occurs with equivalent efficiency in vector transfected and σ3 overexpressing cells (Fig. 2A). As an alternative measure of IKK activity, we determined the capacity of TNFα treatment to mediate p65 Ser536 phosphorylation. In cells expressing σ3, TNFα-induced p65 Ser536 phosphorylation occurs efficiently (Fig. 2B). Consistent with intact IKK activity, we found that σ3 did not impact the steady state levels of IKKβ. We previously demonstrated that IKKβ and NEMO levels are diminished following infection with reovirus strain T3A (13). These data indicate that the mechanism by which σ3 inhibits NF-κB-dependent gene expression is distinct from that observed in the context of T3A infection. Further, these data suggest that reovirus may have evolved to inhibit NF-κB-dependent gene expression in more than one way.

**Fig 2.**
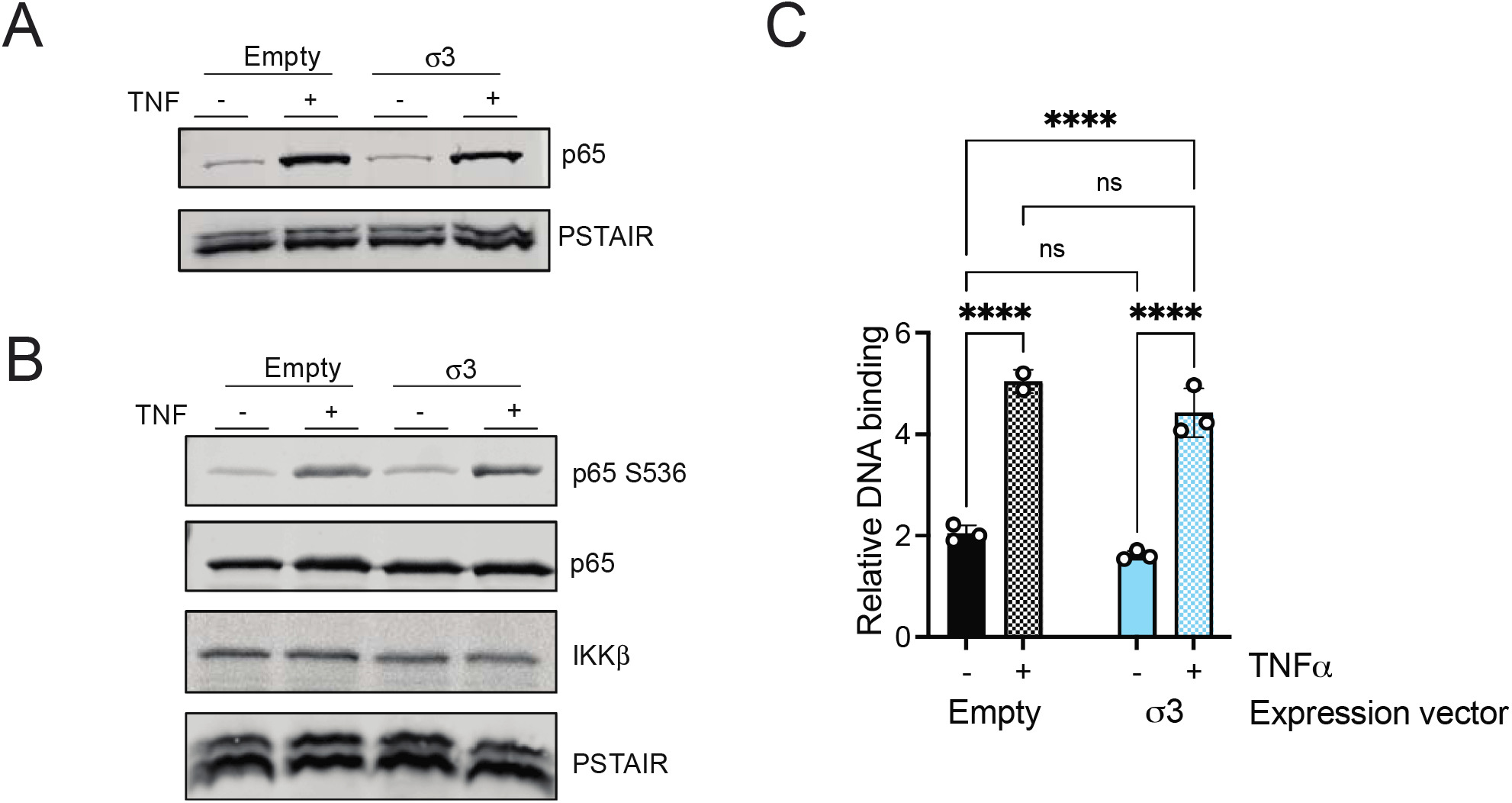
Reovirus protein σ3 does not inhibit upstream NF-κB signaling steps (A) HEK293 cells were transfected with an empty vector or a vector expressing σ3. Following incubation at 37°C for 24 h, cells were treated with 10 ng/ml TNFα and incubated for 1 h. Nuclear extracts were immunoblotted using antiserum specific for p65 or PSTAIR. (B) HEK293 cells were transfected with an empty vector or a vector expressing σ3. Following incubation at 37°C for 24 h, cells were treated with 20 μM proteasome inhibitor PSI for 1 h, then 10 ng/ml TNFα for 30 min. Whole cell extracts were immunoblotted with antisera specific for p65, p65 phosphorylated at Ser536, IKKβ, and PSTAIR. (C) HEK293 cells were transfected with an empty vector or a vector expressing σ3. Following incubation at 37°C for 24 h, cells were treated with 10 ng/ml TNFα for 1 h. Nuclear extracts were assayed for capacity to bind to the NF-κB DNA consensus sequence using an ELISA assay. ****, P < 0.0001 by one-way ANOVA with Tukey’s multiple comparison test. ns indicates not significant.

Data presented thus far indicate that, in σ3 expressing cells, phosphorylated p65 enters the nucleus upon treatment with TNFα, but is unable to induce gene expression. Thus, its nuclear function may be affected. In the nucleus, p65 binds DNAs containing NF-κB binding sites and drives target gene expression by cooperating with transcriptional co-activators. To determine if σ3 influences the DNA binding property of NF-κB, we assessed the capacity of NF-κB to bind its target promoter using a DNA-binding ELISA (Fig. 2C). p65 in lysates of empty vector transfected cells bound to a dsDNA fragment that corresponds to the NF-κB consensus sequence following treatment with TNFα. TNFα-induced interaction between p65 and NF-κB consensus DNA sequence was unaffected in lysates of σ3 expressing cells. These data indicate that the DNA binding by TNFα-activated p65 is not diminished by the presence of σ3.

### σ3 can inhibit NF-κB activated by multiple agonists

In reovirus-infected cells, reovirus genomic dsRNA activates the innate immune response via RIG-I and MDA5 (10, 12). To determine if σ3 inhibits NF-κB-dependent gene expression downstream of RIG-I and MDA5, we co-transfected HEK293 cells with a vector expressing σ3 along with NF-κB-dependent firefly luciferase reporter. We then transfected cells with genomic dsRNA extracted from reovirus particles. In empty vector transfected cells, dsRNA potently stimulated NF-κB-driven reporter gene expression. Similar to our findings with TNFα, we found that cells expressing σ3 showed lower dsRNA-induced NF-κB-dependent gene expression (Fig. 3A). σ3 is a dsRNA binding protein (19, 20). Thus, one way in which σ3 inhibits dsRNA-induced NF-κB activation could be through sequestration of the transfected dsRNA or by preventing its interaction with a sensor. To evaluate this possibility, we compared the capacity of wild-type and dsRNA binding mutant (K293T) σ3 to impact dsRNA-induced NF-κB gene expression (21). We found that wild-type and mutant σ3 equivalently block NF-κB-dependent luciferase expression by dsRNA (Fig. 3A). The σ3 K293T mutant also remained capable of blocking NF-κB activated by TNFα (Fig. 3A). Measurement of steady state levels of wild-type and mutant σ3 indicate that the mutant is capable of blocking NF-κB despite a slightly lower level of expression (Fig. 3B). These data indicate that σ3 can inhibit NF-κB signaling initiated by at least two distinct agonists. Further, our work provides evidence that the previously recognized dsRNA binding region of σ3 does not impact the property of σ3 to inhibit NF-κB.

**Fig 3.**
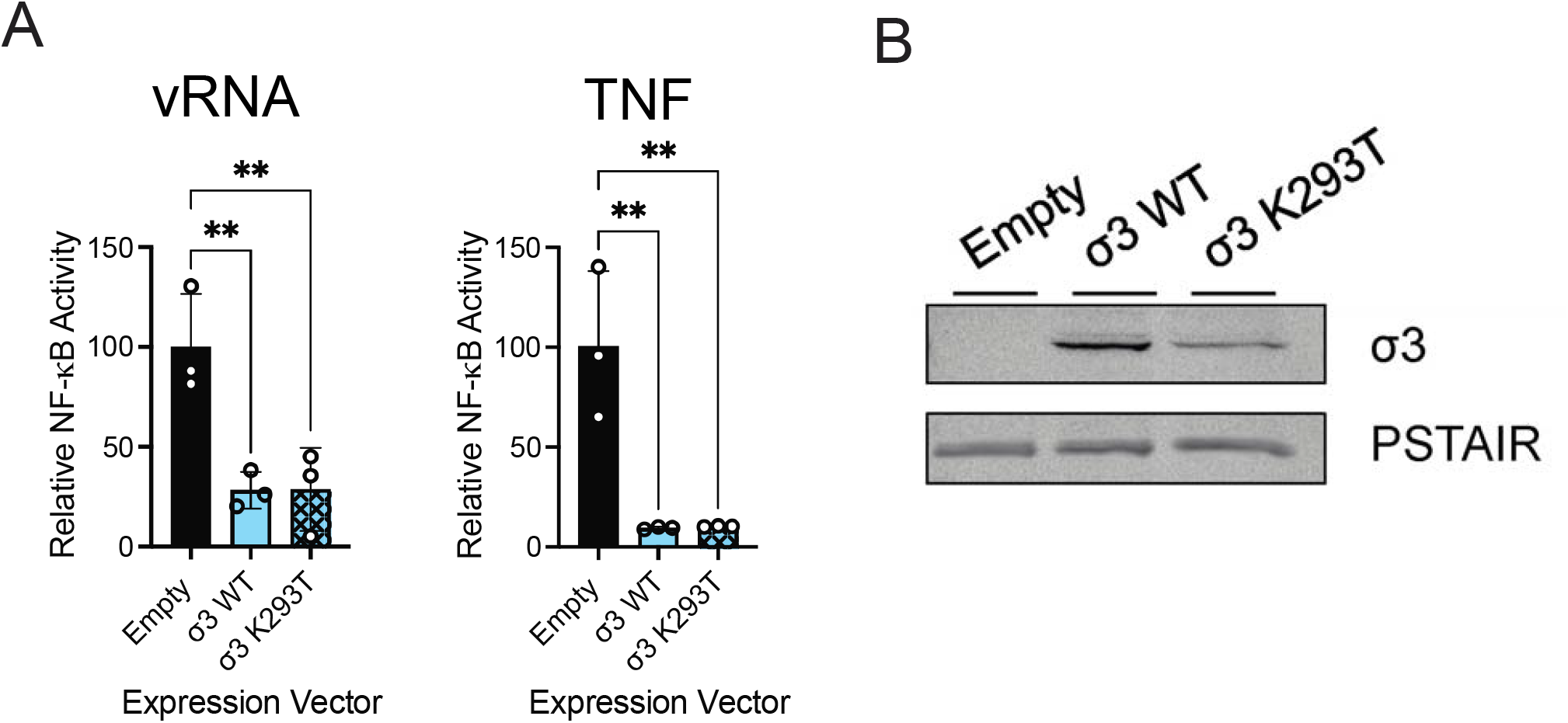
dsRNA binding domain of σ3 is not required for inhibition of NF-κB-dependent gene expression. (A) HEK293 cells were transfected with an empty vector, or with vectors expressing either σ3 WT or σ3 K293T, an NF-κB-driven Firefly luciferase reporter, and a control Renilla luciferase reporter. Following incubation at 37°C for 24 h, HEK293 cells were treated with 100 ng vRNA for 24 h (left panel), or 10 ng/ml TNFα and incubated at for 7 h (right panel). The ratio of Firefly to Renilla luciferase activity (Relative NF-κB activity) was quantified. Relative NF-κB activity in empty vector transfected cells was set to 100%. Activity for each independent transfection and treatment, the mean value and SD are shown. **, P<0.01 by one-way ANOVA with Dunnett’s multiple comparison test. (B) HEK293 cells were transfected with vectors expressing σ3 WT or σ3 K293T. Following an additional incubation at 37°C for 24 h, whole cell extracts were immunoblotted using antiserum specific for reovirus and PSTAIR.

### σ3 selectively inhibits the expression of a subset of NF-κB targets

In experiments completed thus far, we used reporter plasmids to assess the impact of σ3 on NF-κB-dependent gene expression. Promoters of reporter plasmids are much simpler compared to those of endogenous genes. Furthermore, plasmids may undergo a different level of chromatinization compared to endogenous DNA. Thus, the genomic context of a gene may affect its regulation. Toward the goal of assessing the NF-κB inhibitory function of σ3 on endogenous targets, we first confirmed using RT-qPCR that the expression of three known endogenous NF-κB target genes, IκBα, IL-8, and A20 is regulated in an NF-κB-dependent manner in our cells (22-24). The expression of each gene was induced by TNFα and was sensitive to the presence of an IKK inhibitor indicating that each of these genes is NF-κB regulated (Fig. 4A). To determine the impact of σ3 on expression of these three endogenous genes, the capacity of TNFα to stimulate their expression in presence or absence of σ3 was compared. In comparison to empty vector transfected cells, TNFα induced the expression of IL-8 to a lower extent in cells expressing σ3 (Fig. 4B). In contrast, the induction of IκBα and A20 were not significantly affected by the presence of σ3. These data indicate that σ3 only inhibits the expression of some endogenous NF-κB targets.

**Fig 4.**
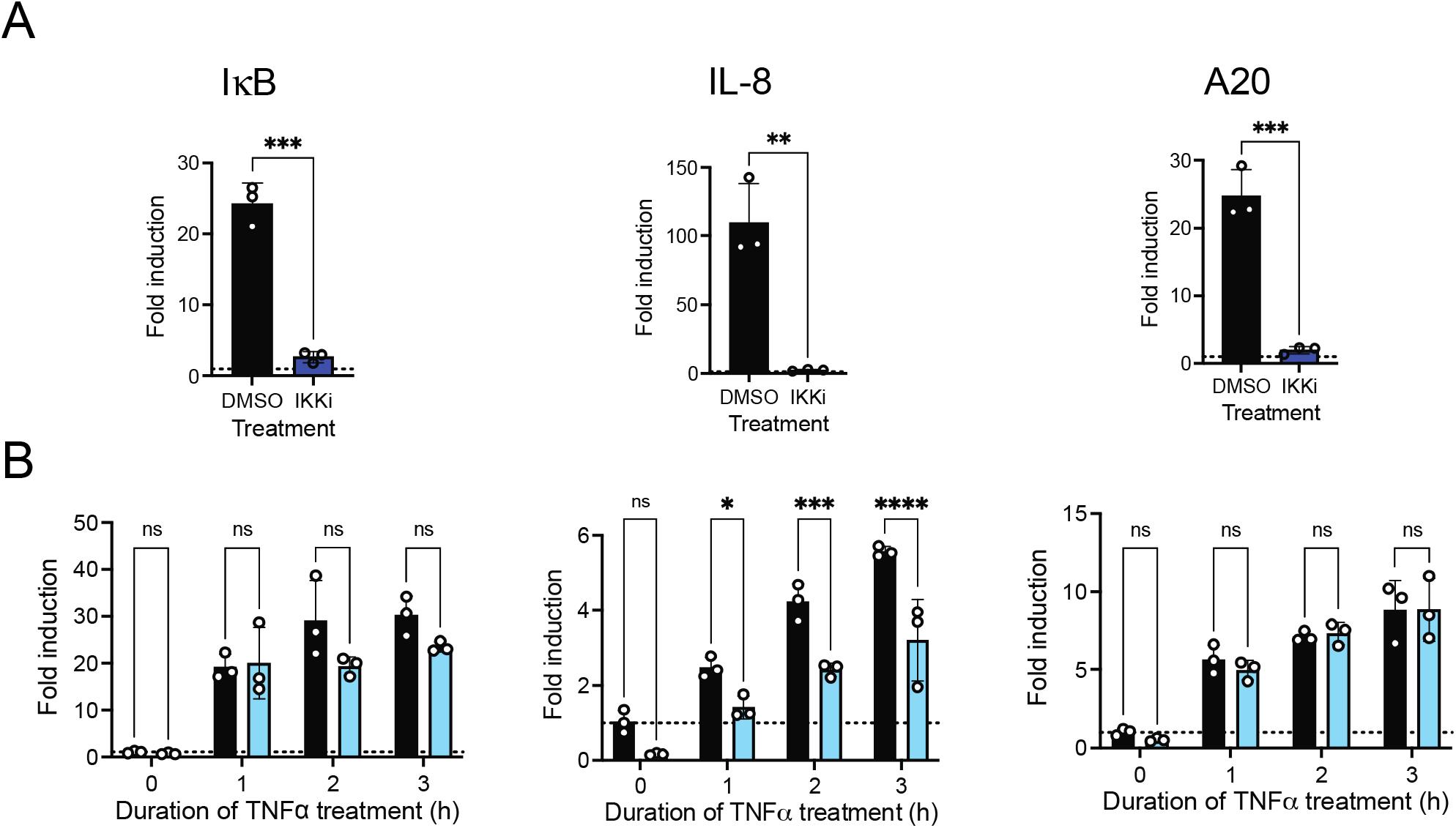
σ3 inhibits expression of a subset of NF-κB target genes. (A) HEK293 cells were treated with DMSO or 5 μM IKK inhibitor BAY-65-1942. Following incubation at 37°C for 1 h, cells were treated with 10 ng/ml TNFα and incubated for 3 h. RNA was extracted from cells and the levels of IL-8, TNFα, and IκBα mRNA relative to GAPDH was measured using qRT-PCR and comparative C_T_ analysis. The ratio of each mRNA relative to GAPDH in control cells in absence of TNFα was set to 1 and is denoted by a dotted line. The ratios for three independent TNFα treatment in DMSO and IKK inhibitor treated cells, the mean and SD are shown. **, P < 0.01, ***, P < 0.001 by Student’s t test in comparison to DMSO treated cells. (B) HEK293 cells were transfected with an empty vector (black bars) or a vector expressing σ3 (blue bars). Following incubation at 37°C for 24 h, cells treated with 10 ng/ml TNFα and incubated for 0, 1, 2, or 3 h. RNA was extracted from cells and the levels of IL-8, TNFα, and IκBα mRNA relative to GAPDH was measured using qRT-PCR and comparative C_T_ analysis. The mean value for three independent transfections and treatments and SD are shown. The ratio of each mRNA relative to GAPDH in control cells in absence of TNFα at 0 h was set to 1 and is denoted by a dotted line.*, P < 0.05, **, P < 0.01 by Student’s t test in comparison to empty vector transfected cells treated with TNFα at the same time interval.

### σ3 inhibits IFN expression through NF-κB inhibition

NF-κB controls the expression of 100s of cellular targets (25). Among genes whose expression is controlled by NF-κB is type I interferon, IFNβ (26). We recently demonstrated that siRNA mediated knockdown of σ3 expression in reovirus-infected cells results in enhancement of IFN expression (27). Our results presented here using ectopic expression of σ3 raise the possibility that σ3 negatively regulates IFNβ expression by repressing NF-κB function. To test this idea, we first determined if IFNβ expression is controlled by NF-κB in our cells. We found that both basal level expression of IFNβ and UV inactivated virus-induced expression of IFNβ is diminished by treatment of cells with an IKK inhibitor indicating that IFNβ expression in our cells is NF-κB-dependent (Fig 5A, B). Expression of σ3 also potently blocked both basal and induced IFNβ expression (Fig 5A, B). These data indicate that the presence of σ3 is sufficient to diminish IFNβ.

**Fig 5.**
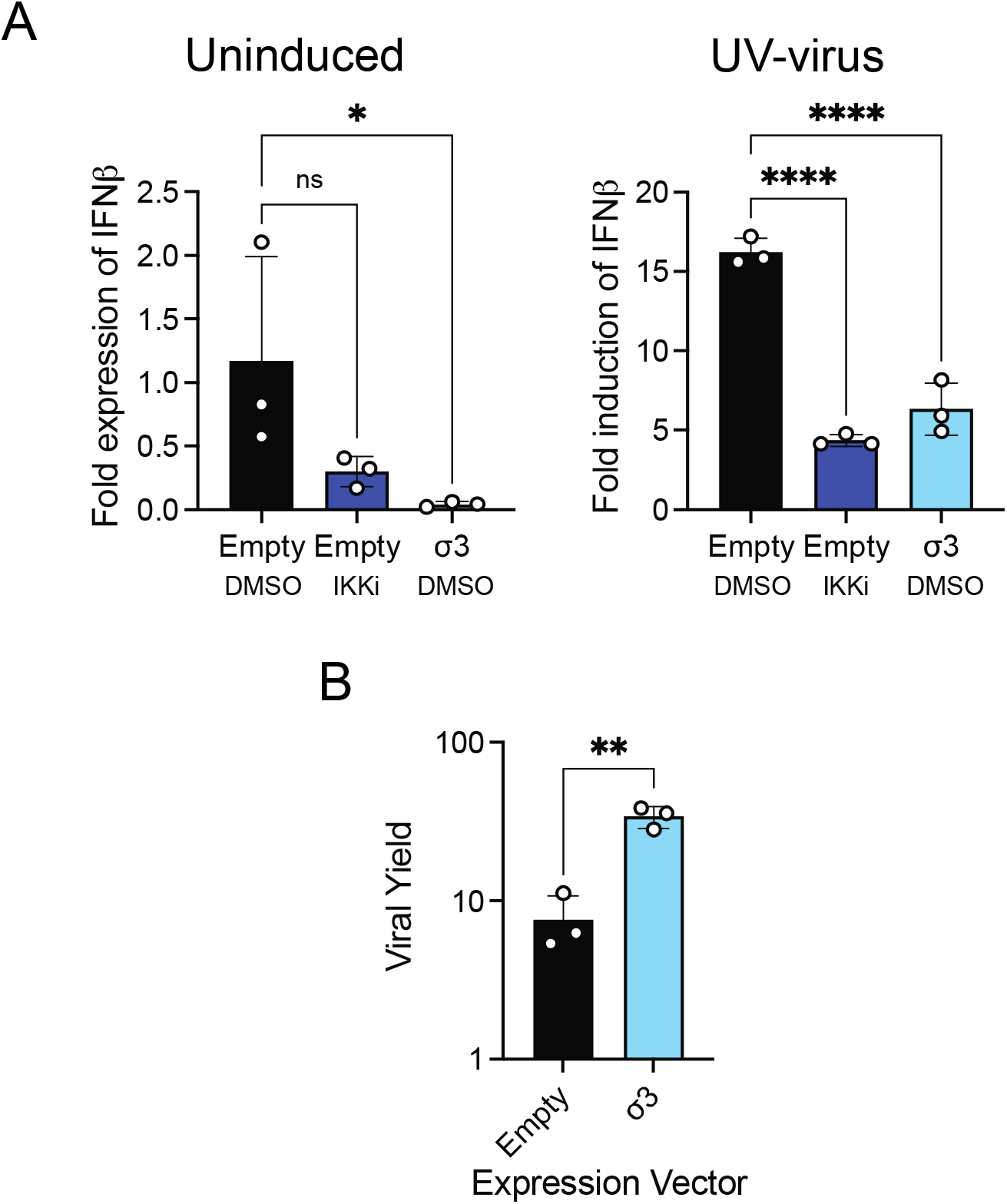
σ3 inhibits IFNβ expression through NF-κB inhibition. HEK293 cells were transfected with an empty vector or a vector expressing tagged σ3. Following incubation at 37°C for 24 h, cells were treated with 0 or 5 μM IKK inhibitor BAY-65-1942. Cells were then absorbed with PBS (mock) (Left panel) or with 10 PFU/cell UV-treated T3D^L^ (Right panel) and incubated at 37°C for 16 h. RNA was extracted from cells and the levels of IFNβ mRNA relative to GAPDH was measured using qRT-PCR and comparative CT analysis. *, P < 0.05, ****, P < 0.0001 by one way ANOVA with Dunnett’s multiple correction test. ns indicates not significant. (B) HEK293 cells were transfected with an empty vector or vector expressing σ3. Following incubation at 37°C for 24 h, the cells were absorbed with 0.1 pfu/cell of reovirus strain T3D^F^. Increase in viral titer over 24 h (viral yield) following infection was measured by plaque assay on L929 cells. **, P < 0.01 by Student’s t test in comparison to empty vector transfected cells.

Because NF-κB controls the expression of antiviral cytokines such as IFNβ, its activation usually results in diminishment of infection (28). Since σ3 inhibits NF-κB activation, we hypothesized that the presence of σ3 generates an environment in the cell that is more favorable to infection. To test this idea, cells transfected with an empty vector or a σ3 expression vector were absorbed with reovirus strain T3D^F^ at an MOI of 0.1 PFU/cell. Viral yield was quantified at 24 h post infection. We found that cells expressing σ3 produced 5-fold more virus than control cells (Fig 5C). These data suggest that σ3 expression likely modifies the cellular environment allowing for greater viral replication.

### σ3 impacts influences the expression of NF-κB target genes during infection

Closely related reovirus strains are known to display differences in a variety of phenotypes (29-31). In previous work we showed that in cells infected with reovirus strain T3A, TNFα fails to induce the expression of NF-κB target genes due to loss of IKKβ and NEMO (13). We found that levels of IKKβ in mock infected and T3D^L^ (a T3D isolate regenerated from plasmids using a T3D isolate from Patrick Lee’s laboratory) are similar. Thus, unlike T3A, T3D^L^ does not diminish IKKβ levels (Fig. 6A). The inability of this virus strain to alter IKKβ levels offered us an opportunity to evaluate the impact of σ3 on NF-κB activity in the context of infection. Based on our experiments reporting that transfection of T3D^F^ σ3 blocks NF-κB signaling, we used reverse genetics to generate a reassortant strain, T3D^L^/T3D^F^S4, that bears a σ3-encoding S4 gene from T3D^F^ in a T3D^L^ genetic background. Similar to T3D^L^, T3D^L^/T3D^F^S4 infection also did not result in the loss of IKKβ (Fig. 6A). In comparison to T3D^L^, the level of σ3 24 h post infection in cells infected with T3D^L^/T3D^F^S4 was higher (Fig. 6A). Under the same conditions, T3D^L^ induced a significantly higher level of NF-κB target genes IL-8 and IFNβ than T3D^L^/T3D^F^S4 (Figure 6B, 6C). The higher level expression of NF-κB targets following infection with T3D^L^ and could be due to better detection of the T3D^L^ genomic material or due to more potent capacity of T3D^L^/T3D^F^S4 to inhibit NF-κB-dependent gene expression. To distinguish between these possibilities, we initiated infection using UV-inactivated viruses. UV-treated reovirus can activate NF-κB and induce IFNβ expression but is not capable of transcribing viral RNAs (10). Remarkably, UV-treated T3D^L^/T3D^F^S4 induced higher (though not statistically different) IFNβ expression than UV-treated T3D^L^ (Fig. 6D). These results indicate that the difference in IFNβ expression by T3D^L^ and T3D^L^/T3D^F^S4 is not due to lower level detection of T3D^L^/T3D^F^S4. Importantly, however, because lower level IFNβ expression for T3D^L^/T3D^F^S4 is only observed when viral replication and gene expression are allowed to continue, these data indicate that a de novo synthesized viral gene product limits IFNβ expression. Since T3D^L^ and T3D^L^/T3D^F^S4 differ only in one gene product, the σ3 protein, these data suggest that the lower level IFNβ expression by T3D^L^/T3D^F^S4 is due to the NF-κB inhibiting function of T3D^F^ derived σ3. Importantly, this work indicates that σ3 properties can impact NF-κB in the context of infection.

**Figure 6.**
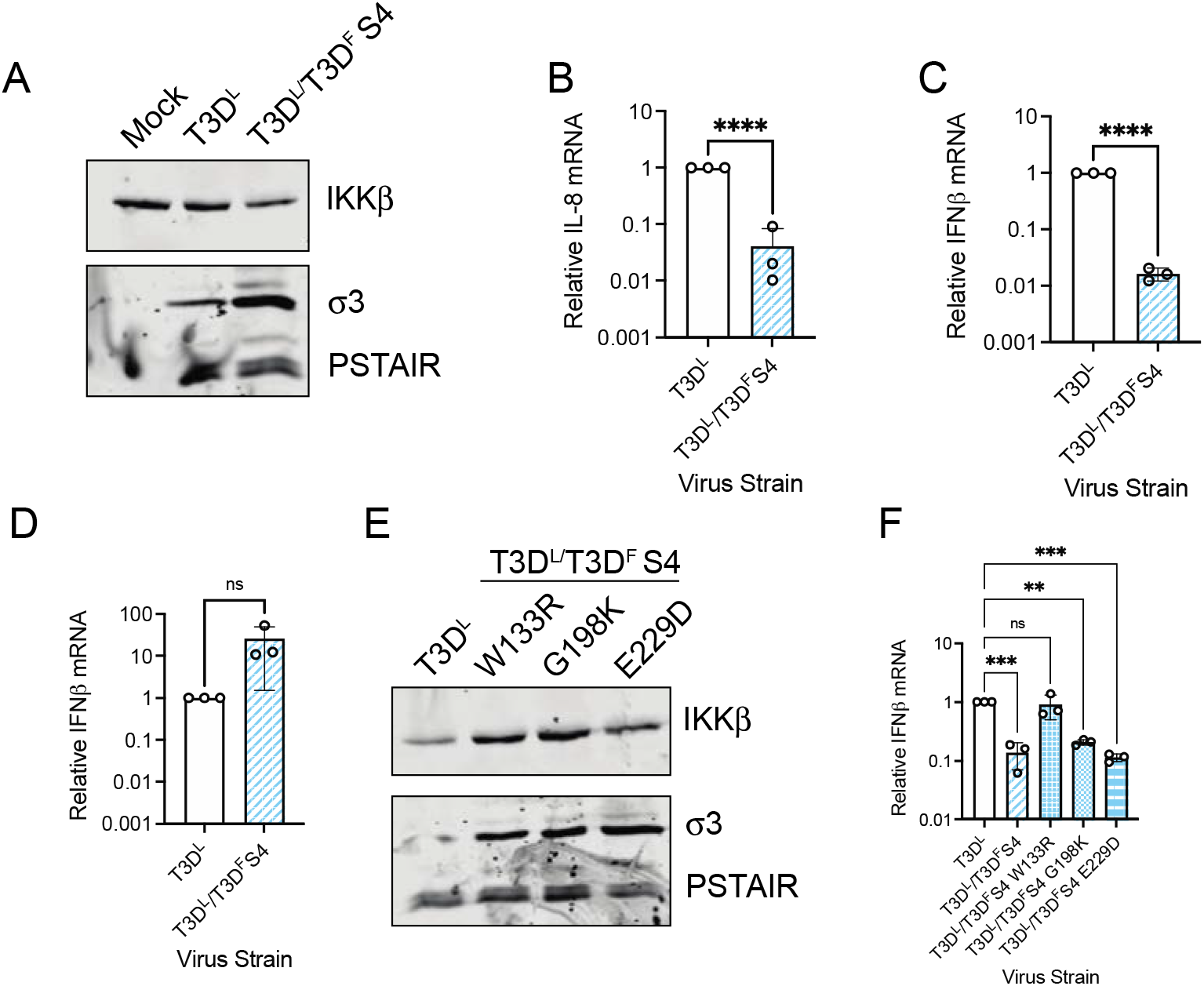
Sigma3 properties influence IFNβ expression during reovirus infection. (A) HEK293 cells were mock infected, or infected with 10 PFU/cell of T3D^L^ or T3D^L^/T3D^F^ S4 for 24 h. Whole cell extracts were immunoblotted using antisera specific for IKKβ, σ3, and PSTAIR. (B,C) HEK293 cells were infected with 10 PFU/cell of T3D^L^ or T3D^L^/T3D^F^ S4 for 24 h. RNA was extracted from cells and the levels of IL-8 (B) and IFNβ (C) relative to GAPDH was measured using qRT-PCR and comparative C_T_ analysis. The mean value for three independent transfections and treatments and SD are shown. The ratio of each mRNA relative to GAPDH in T3D^L^ infected cells was set to 1. ****, P < 0.0001 by student’s t test. (D) HEK293 cells were infected with 10 PFU/cell of UV treated T3D^L^ or UV-treated T3D^L^/T3D^F^ S4 for 12 h. RNA was extracted from cells and the levels IFNβ relative to GAPDH was measured using qRT-PCR and comparative C_T_ analysis. The mean value for three independent transfections and treatments and SD are shown. The ratio of each mRNA relative to GAPDH in UV-T3D^L^ infected cells was set to 1. ****, P < 0.0001 by student’s t test. (E) HEK293 cells were mock infected, or infected with 10 PFU/cell of T3D^L^ or each indicated T3D^L^/T3D^F^ S4 mutant.,Whole cell extracts were immunoblotted using antisera specific for IKKβ, σ3, and PSTAIR. (F) HEK293 cells were infected with 10 PFU/cell of T3D^L^, T3D^L^/T3D^F^ S4 or the indicated T3D^L^/T3D^F^ S4 mutant for 24 h. RNA was extracted from cells and the levels of and IFNβ relative to GAPDH was measured using qRT-PCR and comparative C_T_ analysis. The mean value for three independent transfections and treatments and SD are shown. The ratio of each mRNA relative to GAPDH in T3D^L^ infected cells was set to 1. **, P < 0.01; ***, P < 0.001 by one way ANOVA with Dunnett’s multiple comparison test. ns indicates not significant.

T3D^F^ and T3D^L^ σ3 proteins differ only at three amino acid residues (32). To define the genetic determinants within σ3 that control NF-κB signaling, we generated three T3D^L^ viruses that encode T3D^F^ σ3 incorporating T3D^L^ amino acids at each T3D^F^-T3D^L^ polymorphic residue. The levels of IKKβ in cells infected with each of these viruses was at least as high as that observed for infection with T3D^L^ (Fig. 6E). 24 h following infection, each of these point mutant viruses displayed equivalent expression of σ3. This level appeared higher than that observed for T3D^L^. We found that that virus with W133R change in T3D^F^ σ3 induced IFNβ expression equivalent to T3D^L^ (Fig. 6F). In contrast, G198K and E229D changes in T3D^F^ σ3 induced lower level IFNβ expression similar to T3D^L^/T3D^F^S4. These data indicate that the polymorphic differences at amino acid 133 in σ3 impact the capacity of this protein to inhibit NF-κB. Moreover, because steady state levels of σ3 did not correlate with the expression of IFNβ by these viruses, the data indicate that difference in the properties of σ3 and likely not their amounts in infected cell influence the capacity of σ3 to block NF-κB. Together, our results indicate that σ3 impacts NF-κB activity and consequent IFNβ expression even in the context of infection. This work identifies σ3 as a viral antagonist of the innate immune system.

## DISCUSSION

In this study, we sought to identify the viral requirements for inhibition of NF-κB. We found that the reovirus σ3 protein is sufficient to inhibit TNFα- or dsRNA-induced NF-κB-dependent gene expression (Fig. 1 and 3). Following infection, reovirus inhibits NF-κB by causing a loss of the IKK complex, thereby inhibiting downstream signaling steps. However, nuclear accumulation and phosphorylation of p65 are not inhibited in cells expressing σ3 suggesting that IKKs are fully functional (Fig. 2). Thus, σ3 inhibits NF-κB via a different mechanism than what has been described previously (13). We found that σ3 inhibits the expression of some but not all NF-κB target genes (Fig. 4). The type I IFN, IFNβ is one NF-κB target whose expression is inhibited by the presence of σ3 (Fig. 5). We present data indicating σ3 expression generates an environment in the cell that is more conducive to viral replication (Fig 5.). Our studies also demonstrate that σ3 properties control differences in the expression of NF-κB target genes in cells infected with reovirus variants. These studies uncover a new role for σ3 in antagonizing the host innate immune response.

Despite the multiple mechanisms of NF-κB inhibition, previous work has also shown that reovirus induces NF-κB activity (16, 18). The discrepancy here likely lies in the timing. De novo expression of reovirus gene products is dispensable for reovirus-induced stimulation of NF-κB activity (33). Thus, NF-κB activation in reovirus-infected cells is mediated by components of the incoming virions and occurs prior to viral gene expression. The current model is that genomic material within incoming virions is detected in the cytoplasm by the RIG-I like receptors (RLRs)(34). RLR activation signals through MAVS and leads to the activation of NF-κB. Inhibition of NF-κB, on the other hand, requires viral gene expression (13). Thus early events lead to temporary activation. However, once new viral gene products such as σ3 are made, NF-κB-dependent gene expression is shut off.

There is a strong selective pressure for viruses to antagonize immune pathways which will lead to diminished viral replication. Here, in conjunction with previous studies, we show that reovirus inhibits NF-κB signaling via multiple mechanisms. Distinct from our previous work, we show here that σ3 expression inhibits NF-κB-dependent gene expression, but not through a loss of the IKK complex (13). Thus p65 accumulates in the nucleus but is unable to induce gene expression. We propose two likely hypotheses pertaining to how σ3 inhibits NF-κB-dependent gene expression. First, σ3 alters post-translational modifications of p65. We show here that σ3 does not inhibit p65 Ser536 phosphorylation, however our data cannot rule out the possibility that other modifications of p65 are affected by σ3 (35). These modifications are required for downstream interactions with transcriptional co-activators and chromatin modifying enzymes (36). Different promoters require distinct combinations of transcription factors and co-activators. Such a requirement could account for the inhibition of only a subset of NF-κB target genes (36). Second, although σ3 does not affect the binding of the NF-κB consensus oligo in vitro, it may inhibit DNA binding by p65 to specific promoters. This effect may be direct because σ3 in infected and transfected cells may localize to the nucleus (37, 38), through other cellular proteins, or by changing the chromatin landscape. Though our work identified a residue in σ3 that is important for its NF-κB inhibiting activity, how this residue in σ3 impact NF-κB remains to be defined. In addition to the possibilities described above, amino acid changes in σ3 also affect its function by changing its subcellular localization (38).

Our recent studies also demonstrate that knockdown of σ3 expression enhances IFNβ expression in the context of infection (27). Because σ3 is an essential protein and has multiple activities including in assembling infectious particles, binding dsRNA, controlling translation, and antagonizing PKR and OAS-RNAseL pathways (20, 38-41), whether the effect of σ3 knockdown on IFNβ expression is direct or indirect is hard to decipher. Our studies presented here using ectopically expressed σ3 support a more direct role for σ3 in affecting IFNβ expression through modulating the nuclear function of NF-κB. We also demonstrate that in reovirus infected cells, properties of de novo synthesized σ3 impact the expression of NF-κB target genes. Curiously, polymorphic differences in σ3 between T3D^L^ and T3D^F^ (respectively referred to as T3D-PL and T3D-TD in these studies) have recently been demonstrated to regulate host gene expression (30). T3D^L^ σ3 contributes to the greater capacity of T3D^L^ to express RIG-I- and IFN-independent genes in comparison to T3D^F^. Based on the analysis of promoters of these genes, the authors speculated that RIG-I and IFN-independent genes are regulated in an NF-κB dependent manner. Our studies here demonstrating that T3D^L^/T3D^F^ S4 induces lower level expression of NF-κB target genes than parental T3D^L^ supports this idea. Moreover, data presented here indicate that differences in gene expression correlate with the degree with which NF-κB is inhibited. Plaque size difference in T3D^L^ and T3D^F^ are also controlled by σ3 polymorphisms. The large plaque size of T3D^L^ is diminished upon introduction of T3D^F^ S4 (in T3D^L^/T3D^F^ S4) and then restored by introduction of a single T3D^L^ σ3 residue at amino acid 133 (in T3D^L^/T3D^F^ S4 W133R) (29). Remarkably, plaque size differences shown in that study correlate with the capacity of these reovirus strains to mediate expression of NF-κB target genes (shown in our studies presented here). Whether this relationship is coincidental or reflects a role for the expression of NF-κB target genes in controlling replication or cell to cell spread so as to control plaque size remains to be examined.

Our studies presented here indicate that this impact of σ3 on IFNβ expression is at least in part via the inhibition of NF-kB dependent gene expression. σ3 is thought to enhance translation in cells by preventing PKR activation (19, 20). Two other reoviral proteins antagonize the innate immune response. IRF3 is sequestered in viral factories via the μNS protein limiting IFN expression (42). IRF9 is sequestered in the nucleus in μ2 protein dependent manner inhibiting ISG expression (43). Together, these studies suggest that at least three different reovirus proteins that serve an essential function in the reovirus replication cycle have evolved an additional activity that helps the virus antagonize the host innate immune response.

## MATERIALS AND METHODS

### Cells

HEK293 cells (obtained from M. Marketon’s laboratory) were maintained in Dulbecco’s modified essential medium (DMEM) (Lonza) supplemented with 10% fetal bovine serum (FBS) and 2 mM L-glutamine. Murine L929 cells (ATCC CCL-1) were maintained in Eagle’s minimal essential medium (MEM) (Lonza) supplemented with 10% fetal bovine serum (FBS) and 2mM L-glutamine. Spinner-adapted L929 cells (obtained from T. Dermody’s laboratory) were maintained in Joklik’s MEM (Lonza) supplemented to contain 5% FBS, 2 mM L-glutamine, 100 U/ml of penicillin, 100 µg/ml of streptomycin, and 25 ng/ml of amphotericin B. Spinner-adapted L929 cells were used for cultivating and purifying viruses and for plaque assays.

### Plasmids

Expression vectors for each reovirus T3D^F^ open reading frame in pCAG vector were obtained from the Dermody laboratory. pCMV-T3DF S4 was also obtained from the Dermody laboratory. Plasmids to generate T3D^L^ by reverse genetics were obtained from Takeshi Kobayashi (32). Reverse genetics plasmids expressing T3D^F^ S4 point mutants were obtained from Maya Shmulevitz (29). Firefly and renilla luciferase reporter plasmids are from Promega.

### Reagents and antibodies

The IKK inhibitor, described by Bayer (BAY-45-1962), was used at a concentration of 10 µM. TNFα, purchased from Sigma-Aldrich (T0157), was used at indicated concentrations. Antibodies against p65 were purchased from Santa Cruz Biotechnology (sc-372), those against p65 Ser536 and IKKβ were purchased from Cell signaling (93H1 and D30C6). Anti-PSTAIR monoclonal antibody was purchased from Sigma (P7962). Antisera against reovirus have been previously described (44).

### Viruses

A laboratory stock of T3D^F^ was used generated by plasmid based reverse genetics (45). Infectious viral particles were purified by Vertrel XF extraction and CsCl gradient centrifugation. T3D^L^, T3D^L^/T3D^F^ S4 and its point mutants were generated by plasmid based reverse genetics. Viral titers were determined by a plaque assay using spinner-adapted L929 cell with chymotrypsin in the agar overlay. UV-inactivated virus was generated by using a UV cross-linker (CL-1000 UV cross-linker; UVP). Virus diluted in phosphate-buffered saline (PBS) was placed into a 60-mm tissue culture dish and irradiated with short-wave (254-nm) UV at a distance of 10 cm for 1 min at 120,000 µJ/cm^2^.

### Plasmid transfections

All transfections were done in nearly confluent monolayers of HEK293 cells using 3 µl Lipofectamine 2000/µg plasmid. 12 well plates were transfected with either 1 μg of empty vector or 1 μg of σ3 expression vector. 24 well plates were transfected with either 0.5 μg of empty vector or 0.5 μg of σ3 expression vector. For luciferase assays, 96 well plates were transfected with 0.1 µg/well vectors expressing each viral protein and 0.07 µg/well of an NF-κB or IRF3 reporter plasmid, which expresses firefly luciferase under NF-κB control (pNF-κB-Luc) or IRF3 control (p55-luc), and 0.03 µg/well of control plasmid pRenilla-Luc, which expresses Renilla luciferase constitutively.

### Luciferase assays

At 24 h post-transfection, cells were treated with TNFα for 7 h or transfected with dsRNA for 24 h. Luciferase activity in the cultures was quantified using the Dual-Luciferase Assay Kit (Promega) according to the manufacturer’s instructions.

### RT-qPCR

RNA was extracted using the Aurum Total RNA Mini Kit (Bio-Rad). For RT-qPCR, 0.5 to 2_μg of RNA was reverse transcribed with the high-capacity cDNA RT kit (Applied Biosystems), using random hexamers. cDNA was subjected to PCR using SYBR Select Master Mix using gene specific primers (Applied Biosystems). Fold increases in gene expression with respect to control samples (indicated in each figure legend) were measured using the ΔΔ*C*_*T*_ method (46). Calculations for determining ΔΔ*C*_*T*_ values and relative levels of gene expression were performed as follows: fold increase in cellular gene expression (with respect to glyceraldehyde-3-phosphate dehydrogenase [GAPDH] levels) = 2^−[(gene of interest *CT* – GAPDH *CT*)TNFα – (gene of interest *CT* – GAPDH *CT)*control]^

### Preparation of cellular protein extracts

At 24 h post-transfection, cells were treated with TNFα for a variety of times. For preparation of whole-cell lysates, cells were washed in phosphate-buffered saline (PBS) and lysed with 1× RIPA (50 mM Tris [pH 7.5], 50 mM NaCl, 1% TX-100, 1% deoxycholate, 0.1% SDS, and 1 mM EDTA) containing a protease inhibitor cocktail (Roche), 500 µM dithiothreitol (DTT), and 500 µM phenylmethylsulfonyl fluoride (PMSF), followed by centrifugation at 15,000 × *g* at 4°C for 15 min to remove debris. Nuclear extracts were prepared by lysing cells in a hypotonic lysis buffer (10 mM HEPES, 10 mM KCl, 1.5 mM MgCl, 0.5 mM DTT, and 0.5 mM PMSF for 15 min, subsequent addition of 0.5% NP-40, and 10 seconds of vortexing. After centrifugation at 10,000 x g at 4°C for 10 min, nuclear pellet was washed with hypotonic lysis buffer and then resuspended in high-salt nuclear extraction buffer (25% glycerol, 20mM HEPEs, 0.42 M NaCl, 10 mM KCl, 1.5 mM MgCl, 0.5 mM DTT, 0.5 mM PMSF) at 4°C for 1 h. Nuclear extracts were obtained following removal of the insoluble fraction by centrifugation at 12,000 x g at 4°C for 10 min.

### Immunoblot Assay

Protein concentrations were estimated using a DC Protein Assay from Bio-Rad. Equal protein was loaded, and the cell lysates or extracts were resolved by electrophoresis in 10% polyacrylamide gels and transferred to nitrocellulose membranes. Membranes were blocked for at least 1 h in blocking buffer (StartingBlock T20 TBS Blocking Buffer) and incubated with antisera against p65 (1:1,000), p65 p-Ser536 (1:1,000), IKKβ (1:1,000), reovirus (1:5,000), and PSTAIR (1:5,000) at 4°C overnight. Membranes were washed three times for 5 min each with washing buffer (Tris-buffered saline [TBS] containing 0.1% Tween-20) and incubated with a 1:20,000 dilution of Alexa Fluor-conjugated goat anti-rabbit Ig (for p65, p65 p-Ser536, IKKβ, and reovirus) or goat anti-mouse Ig (for PSTAIR) in blocking buffer. Following three washes, membranes were scanned using a ChemiDoc (Bio-Rad) or Licor Odyssey.

### NF-κB DNA Binding Assay

NF-κB activity was measured by a NF-κB p65 Transcription Factor Assay Kit (Cayman) according to the manufacturer’s instructions.

### Statistical analysis

Statistical significance between experimental groups was determined using the unpaired student’s *t*-test function in excel and graphed using Graphpad Prism software. For experiments involving more than two groups, one-way ANOVA with multiple comparisons was used. Statistical analyses for differences in gene expression by RT-qPCR were done on the ΔC_T_ values.

## ACKNOWLEDGEMENTS

We thank members of our laboratory and the Indiana University Virology community for helpful suggestions. We are also grateful to Dr. Karl Boehme for review of our manuscript.

Research reported in this publication was supported by the National Institute of Allergy and Infectious Diseases of the National Institutes of Health under award numbers R01AI110637 (PD). The content is solely the responsibility of the authors and does not necessarily represent the official views of the funders.

